## Supplemental Figures for "Combinatorial effects of multiple genes contribute to beneficial aneuploidy phenotypes"

### Figure S1

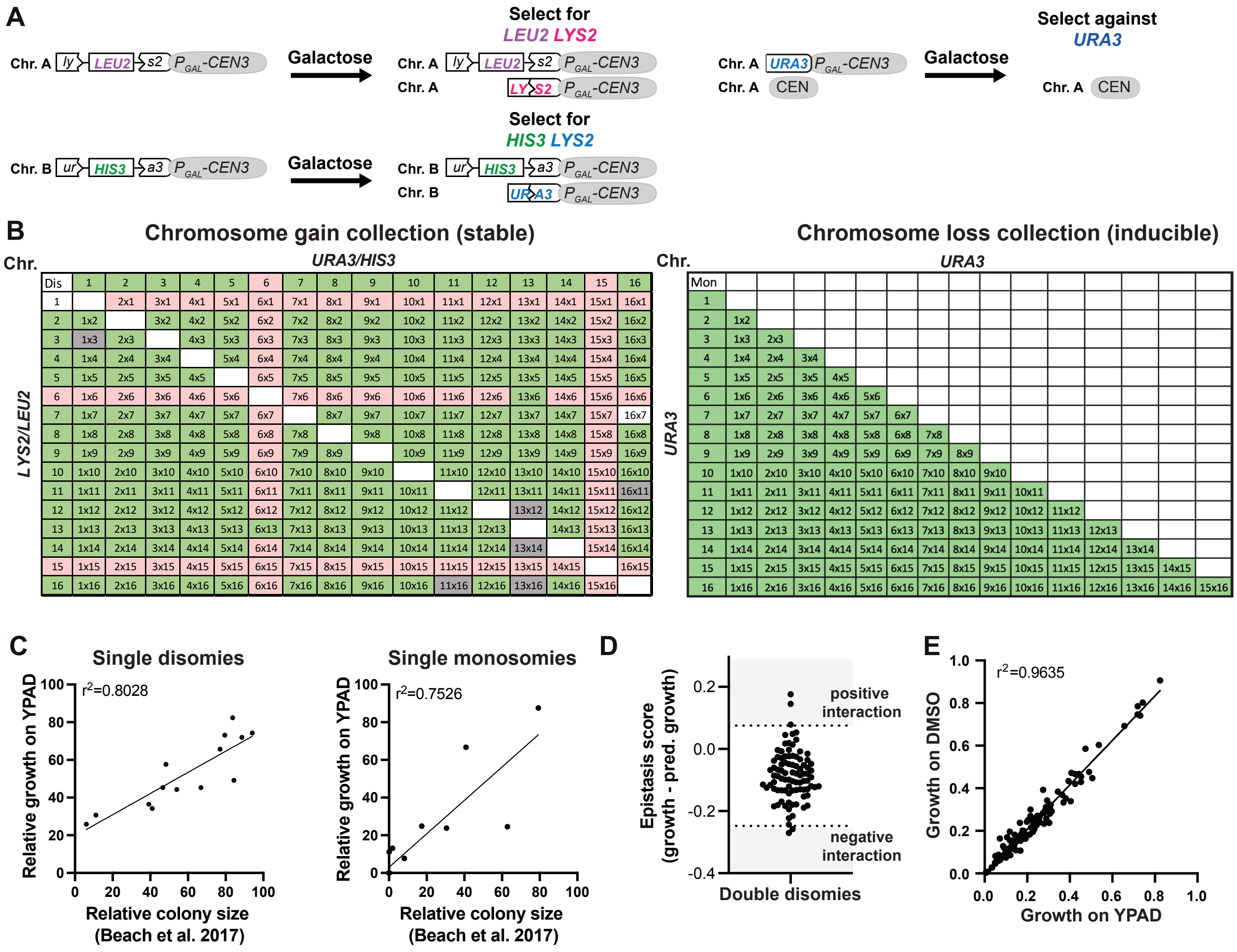


**Figure S1**

(A) Schematic of the galactose-inducible system for engineering specific disomies and monosomies. Strains were grown in medium containing galactose to induce chromosome nondisjunction during mitosis. Monosomy of the desired chromosome was selected for on plates containing 5-Fluoroorotic Acid (5-FOA), selecting against the *URA3* labeled chromosome. Disomy of the desired chromosome was selected for on minimal medium plates lacking leucine, and lysine, or histidine, and uracil, or all 4 amino acids. (B) Overview of the karyotypes represented in the chromosome gain collection (left) and the chromosome loss collection (right). Karyotypes highlighted in green are included in the collection, karyotypes highlighted in grey were excluded from the analysis due to diploidization and those highlighted in red are combinatorial lethal or the euploid parent strain had fitness defects. All possible single monosomies and monosomy combinations are inducible for chromosome loss, but not all are viable. (C) Correlation between growth of the single disomies (left) and monosomies (right) from two independent high throughput spot assays to growth of the single disomies and monosomies from Beach et al. 2017. All strains were normalized to their respective WT controls (disomies: $r^2$ = 0.749, $p < 0.0001$, monosomies: $r^2$ = 0.753, $p < 0.0001$). $r^2$ values are from simple linear regression and p-value are from *F*-tests.. (D) Scatter plot of epistasis scores (measured growth - predicted growth) of the double disomies. Growth predictions are based on the product of the respective single disomies. Kolmogorov-Smirnov was used to test for a normal distribution (KS = 0.05935 and p-value > 0.1). Negative and positive inter chromosomal interactions are defined by z-scores < -2 (-0.248) and > 2 (0.075) and are indicated by dashed lines. (E) Correlation between quantified high throughput spot assays from control plates (YPAD and YPAD + DMSO) ($r^2$ = 0.9634, $p < 0.0001$). $r^2$ values are from simple linear regression and p-value are from *F*-tests.

#### Figure S2

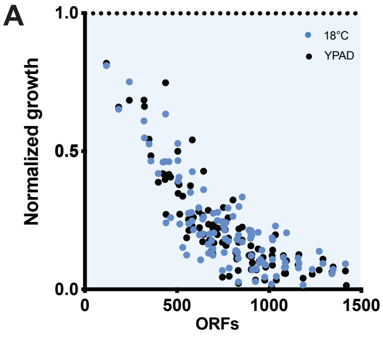


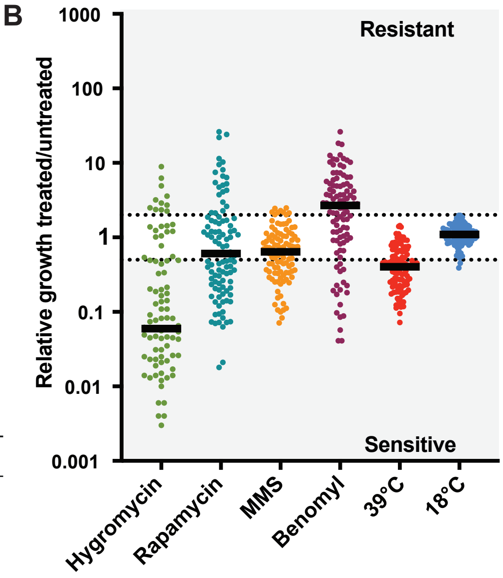


C

Relative growth treated/untreated

| % of strains | 39°C | 18°C | Hygromycin | Rapamycin | MMS | Benomyl |
|---|---|---|---|---|---|---|
| above 2 | 0 | 0 | 11 | 21 | 9 | 59 |
| below 0.5 | 66 | 3 | 78 | 48 | 39 | 17 |

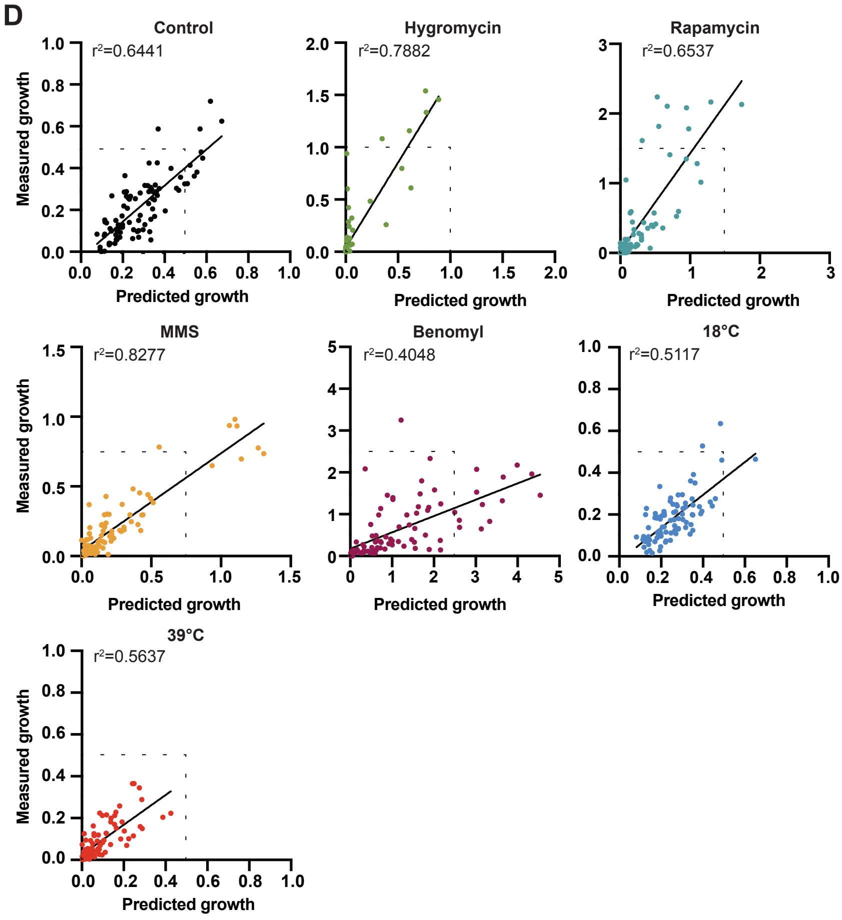


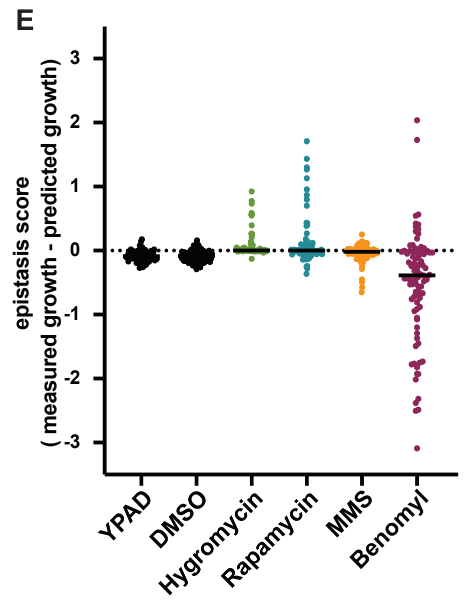


**Figure S2:**

(A) Growth of strains from the chromosome gain collection. Control plates were quantified after 24h (black) and temperature shifted plates after 30 h (39 °C, red) and 48 h (18 °C, blue). All measurements were from two independent experiments and normalized to a haploid WT. (B) Relative growth of the chromosome gain collection under selective conditions normalized to both WT and untreated control plates. Relative growth > 2 is counted as resistant and relative growth < 0.5 as sensitive. (C) Table of the percentage of strains that are resistant or sensitive for each treatment shown in B. (D) Correlations between the growth of double disomies and predicted growth based on single disomies of two independent high-throughput growth assays. Assay conditions: YPAD averaged with YPAD + DMSO (black, $r^2 = 0.6441$, $p < 0.0001$), YPAD + 40 μg/ml hygromycin (green, $r^2 = 0.7882$, $p < 0.0001$), YPAD + 5 nM rapamycin (blue, $r^2 = 0.6537$, $p < 0.0001$), YPAD + 0.02 % MMS (orange, $r^2 = 0.8277$, $p < 0.0001$), YPAD + 17.5 μg/ml benomyl (magenta, $r^2 = 0.4048$, $p < 0.0001$), YPAD under cold stress (18 °C, blue, $r^2 = 0.5117$, $p < 0.0001$) and YPAD under heat stress (39 °C, red, $r^2 = 0.5637$, $p < 0.0001$). The dashed line indicates a perfect match between predicted growth and measured growth. $r^2$ values are from simple linear regression and p-value are from *F*-tests. (E) Epistasis scores under different growth conditions. Negative scores indicate negative genetic interactions and positive scores indicate positive interactions.

**Figure S3**

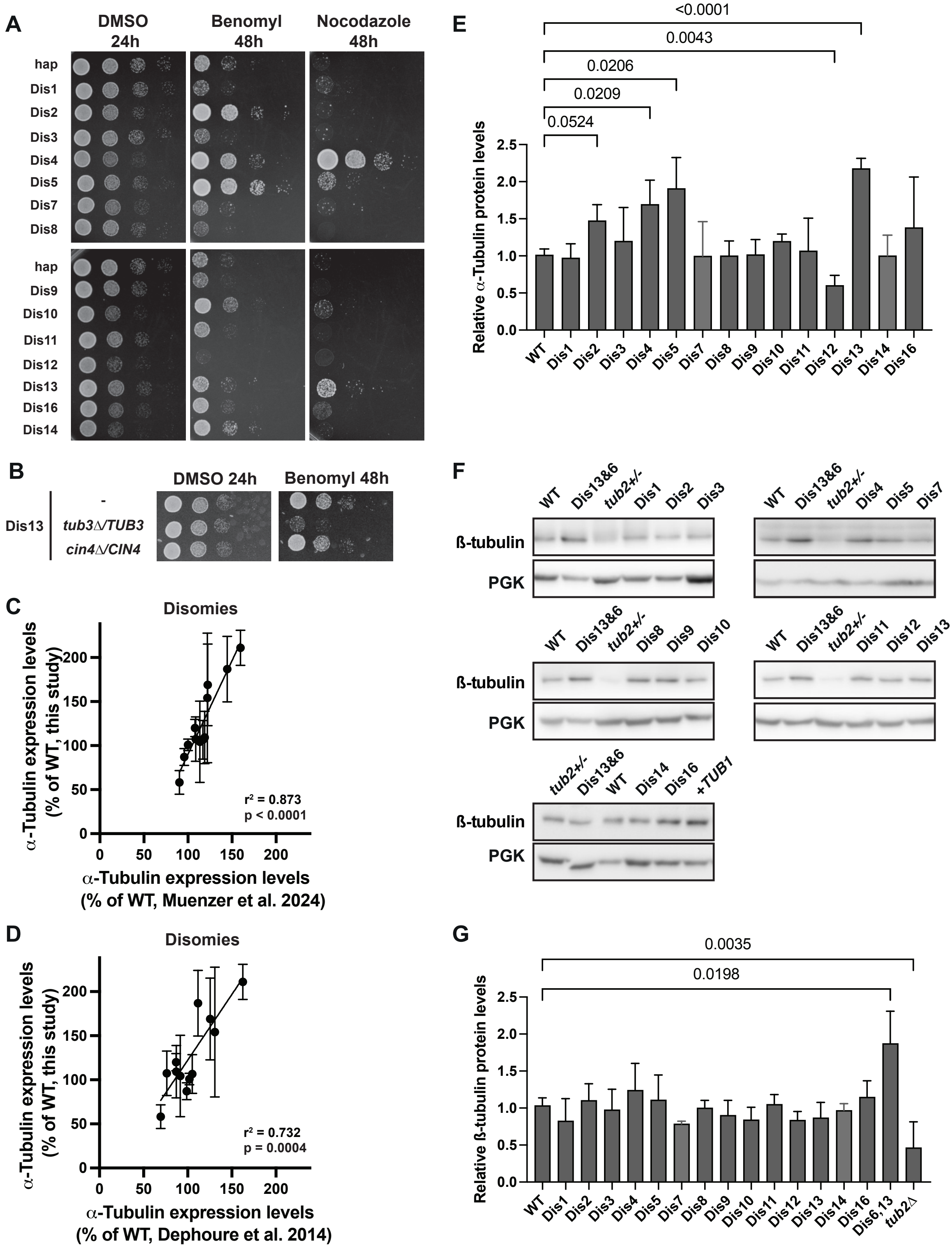
A
DMSO 24h
Benomyl 48h
Nocodazole 48h
hap
Dis1
Dis2
Dis3
Dis4
Dis5
Dis7
Dis8
hap
Dis9
Dis10
Dis11
Dis12
Dis13
Dis16
Dis14
B
DMSO 24h
Benomyl 48h
Dis13
-
tub3∆/TUB3
cin4∆/CIN4
C
Disomies
α-Tubulin expression levels (% of WT, this study)
α-Tubulin expression levels (% of WT, Muenzer et al. 2024)
r² = 0.873
p < 0.0001
D
Disomies
α-Tubulin expression levels (% of WT, this study)
α-Tubulin expression levels (% of WT, Dephoure et al. 2014)
r² = 0.732
p = 0.0004
E
Relative α-Tubulin protein levels
<0.0001
0.0043
0.0206
0.0209
0.0524
WT Dis1 Dis2 Dis3 Dis4 Dis5 Dis7 Dis8 Dis9 Dis10 Dis11 Dis12 Dis13 Dis14 Dis16
F
WT Dis13&6 tub2+/- Dis1 Dis2 Dis3
WT Dis13&6 tub2+/- Dis4 Dis5 Dis7
WT Dis13&6 tub2+/- Dis8 Dis9 Dis10
WT Dis13&6 tub2+/- Dis11 Dis12 Dis13
tub2+/- Dis13&6 WT Dis14 Dis16 +TUB1
ß-tubulin
PGK
G
Relative ß-tubulin protein levels
0.0035
0.0198
WT Dis1 Dis2 Dis3 Dis4 Dis5 Dis7 Dis8 Dis9 Dis10 Dis11 Dis12 Dis13 Dis14 Dis16 Dis6,13 tub2∆

**Fig. S3**

(A) 10-fold serial dilution of all single disomies on agar plates containing DMSO (24h), 25 µg/ml Benomyl, or 10 µg/ml Nocodazole. Disomies 4, 5, 10 and 13 are resistant to both drugs. (B) 10-fold serial dilution of disomy 13, disomy 13 *tub3Δ/TUB3,* and disomy 13 *cin4Δ/CIN4* on YPAD + DMSO (24h) and YPAD + 20 µg/ml benomyl (48h) plates. Full serial dilution image including the WT control is in Supplemental Fig. S7A. (C and D) Correlation between α-tubulin expression levels of the single disomies quantified by western blot (Fig 3B and C) and α-tubulin expression levels of the Torres et al. 2007 single disomies quantified by mass spectrometry. Data from Muenzner et al. 2024 is shown in C and Dephoure et al. 2014 in D. r2 and p-value are calculated from Pearson correlations. (E) Quantification of α-tubulin expression levels of the single disomies quantified by western blot (Fig. 3B, n = 3). Mean and standard deviation are plotted. p-values are calculated using Brown-Forsythe and Welch ANOVA. (F) Western blot analysis of β-tubulin expression levels of all single disomies. The double disomy of chromosomes 6 and 13 and a *tub2Δ/TUB2* diploid strain were used as controls for changes in β-tubulin expression levels. Pgk1 is shown as a loading control. (G) Quantification of ß-tubulin expression levels of the single disomies quantified by western blot (n = 3). Mean and standard deviation are plotted. p-values are calculated using Brown-Forsythe and Welch ANOVA.

#### Figure S4

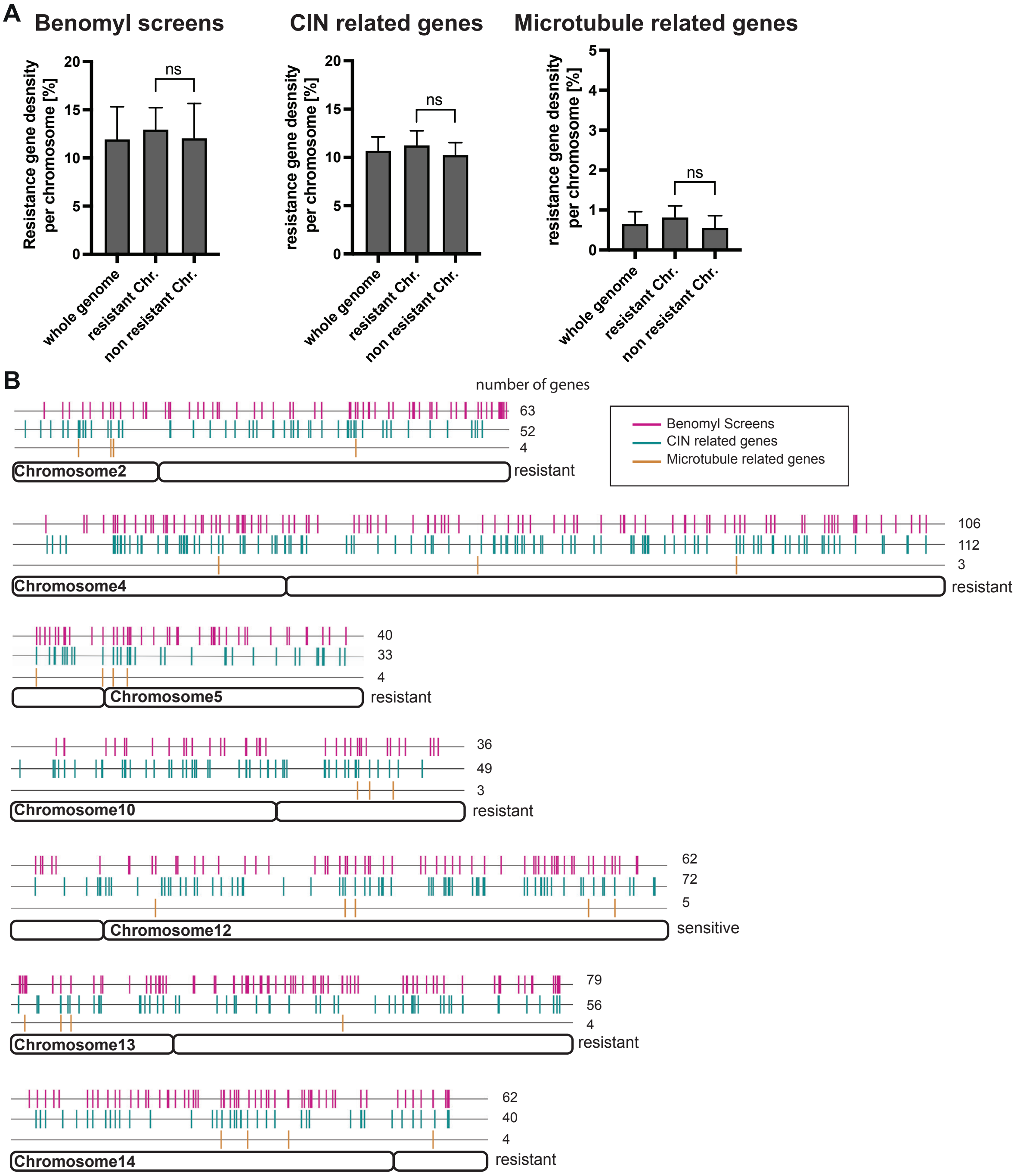


**Fig. S4**

(A) Comparison of the distribution of candidate resistance genes based on different categories of GO terms between resistant chromosomes (2, 4, 5, 10, 13), non-resistant chromosomes, and all chromosomes (whole genome). Candidate resistance gene density was measured relative to all genes on a chromosome for each chromosome individually. Mean and standard deviation for each category are plotted. p-values are calculated using ordinary one-way ANOVA. (B) Distibution of candidate resistance genes for each category on benomyl resistant and sensitive chromosomes.

### Figure S5

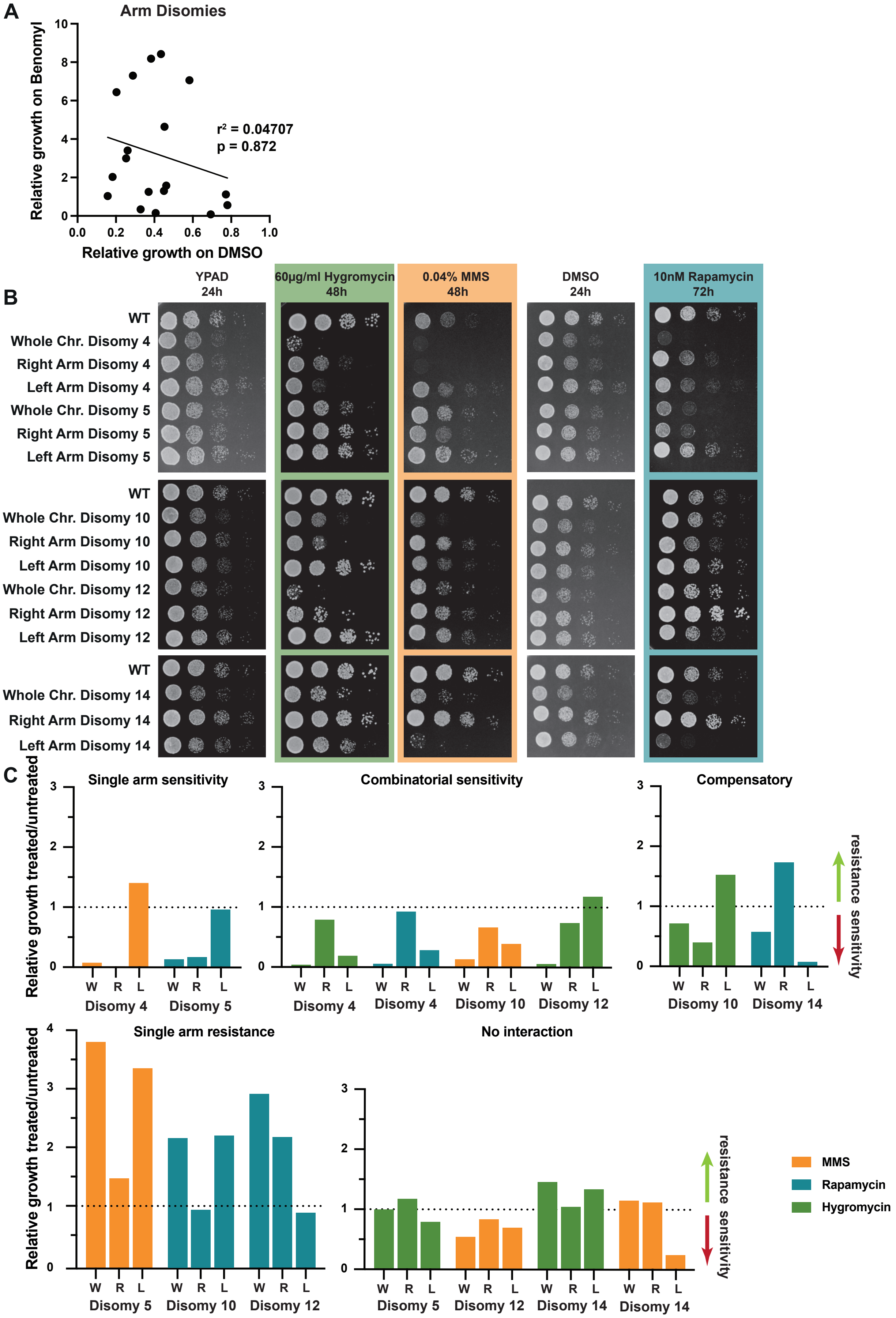
A
Arm Disomies
Relative growth on Benomyl
Relative growth on DMSO
r² = 0.04707
p = 0.872
B
YPAD 24h
60µg/ml Hygromycin 48h
0.04% MMS 48h
DMSO 24h
10nM Rapamycin 72h
WT
Whole Chr. Disomy 4
Right Arm Disomy 4
Left Arm Disomy 4
Whole Chr. Disomy 5
Right Arm Disomy 5
Left Arm Disomy 5
WT
Whole Chr. Disomy 10
Right Arm Disomy 10
Left Arm Disomy 10
Whole Chr. Disomy 12
Right Arm Disomy 12
Left Arm Disomy 12
WT
Whole Chr. Disomy 14
Right Arm Disomy 14
Left Arm Disomy 14
C
Single arm sensitivity
Combinatorial sensitivity
Compensatory
Relative growth treated/untreated
W R L
Disomy 4
Disomy 5
Disomy 10
Disomy 12
Disomy 14
resistance sensitivity
Single arm resistance
No interaction
MMS
Rapamycin
Hygromycin

**Figure S5**

(A) Lack of correlation between growth of the arm level disomies on control (YPAD + DMSO) and treated plates (YPAD + 25 µg/ml benomyl) (r2 = 0.04707, p-value = 0.872). $r^2$ values are from simple linear regression and p-value are from *F*-tests. Quantification of Figure 4D. (B) 10-fold serial dilutions of whole chromosome and single arm aneuploidies on agar plates with YPAD, YPAD + DMSO, YPAD + 60 µg/ml hygromycin (green), YPAD + 0.04 % MMS (orange), or YPAD + 10 nM rapamycin (cyan). Timing of drug condition imaging was adjusted to the strength of the drug condition (hygromycin and MMS 48h, rapamycin 72h). (C) Quantifications of 10-fold serial dilutions from B. On control plates, the 3rd dilution was quantified and on treated plates, the 2nd dilution. Plate growth was first normalized to the haploid WT and then normalized to the untreated control. Each whole chromosome disomy (W) and the corresponding arm level disomies (L = left arm disomy, R = right arm disomy) was assigned to one of five categories: single arm sensitivity – one arm is treatment sensitive, combinatorial sensitive – both chromosome arms contribute to the whole chromosome sensitivity, compensatory sensitivity – a resistant arm can compensate for the sensitivity of a sensitive arm, single arm resistance – a single arm contributes to treatment resistance, and no interaction – whole chromosome disomy and single arm disomies are unaffected by the treatment.

Figure S6

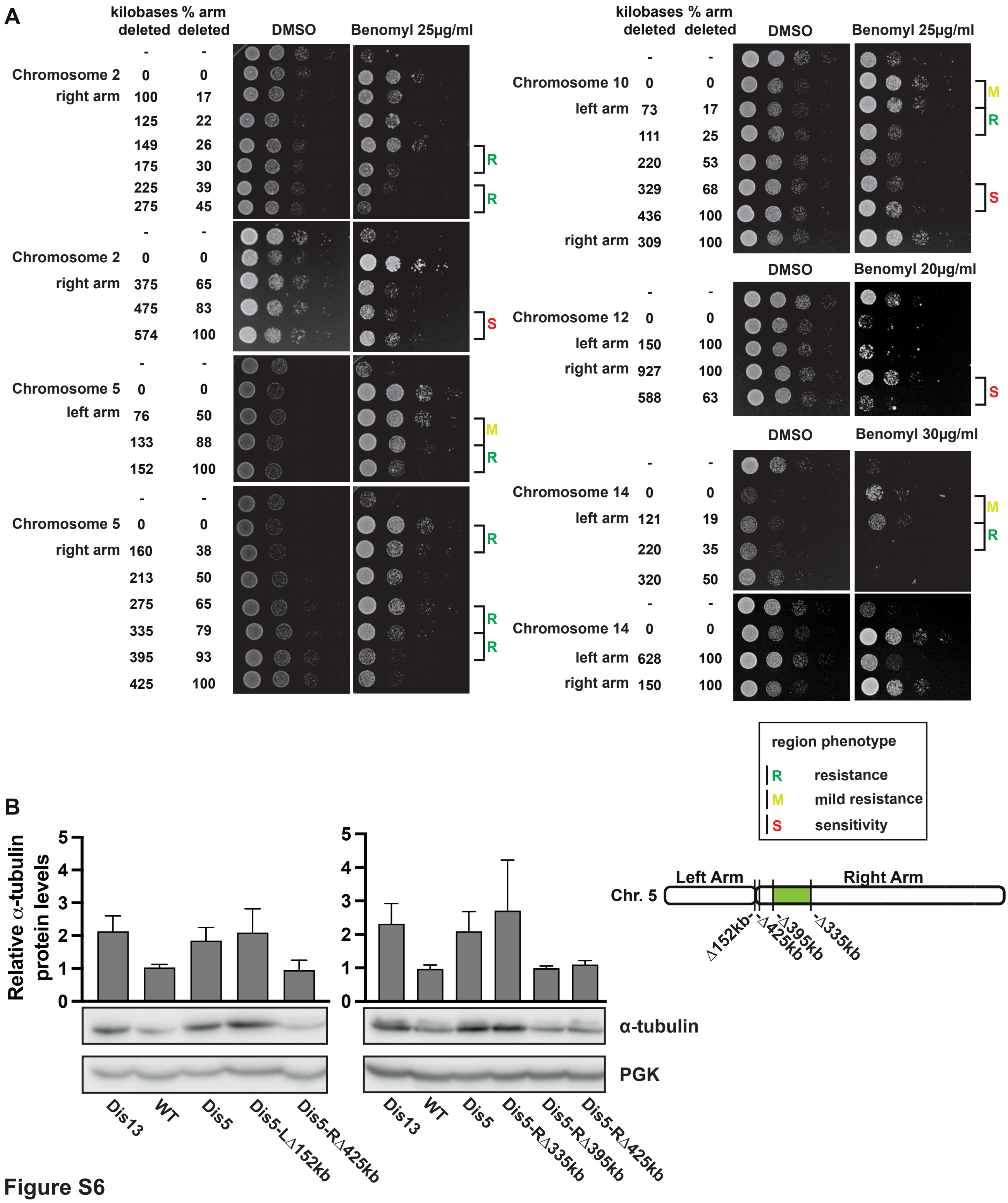


**Figure S6**

(A) 10-fold serial dilutions of partial chromosome arm deletions for chromosomes 2, 4, 5, 10, 12 and 14. Control plates (YPAD + DMSO) were imaged after 24h, and treated plates (YPAD + benomyl) after 48h. (B) Quantification of α-tubulin expression levels of the single disomies quantified by western blot (n = 3). Dis13 is a control for elevated α-tubulin levels. Mean and standard deviation are plotted. Quantified bands were normalazized to the Ponceau stained membrane and the euploid WT. The diagram on the right shows the location of the cuts on the right and left arms of chromosome 5. The region leading to α-tubulin overexpression is in green.

### Figure S7

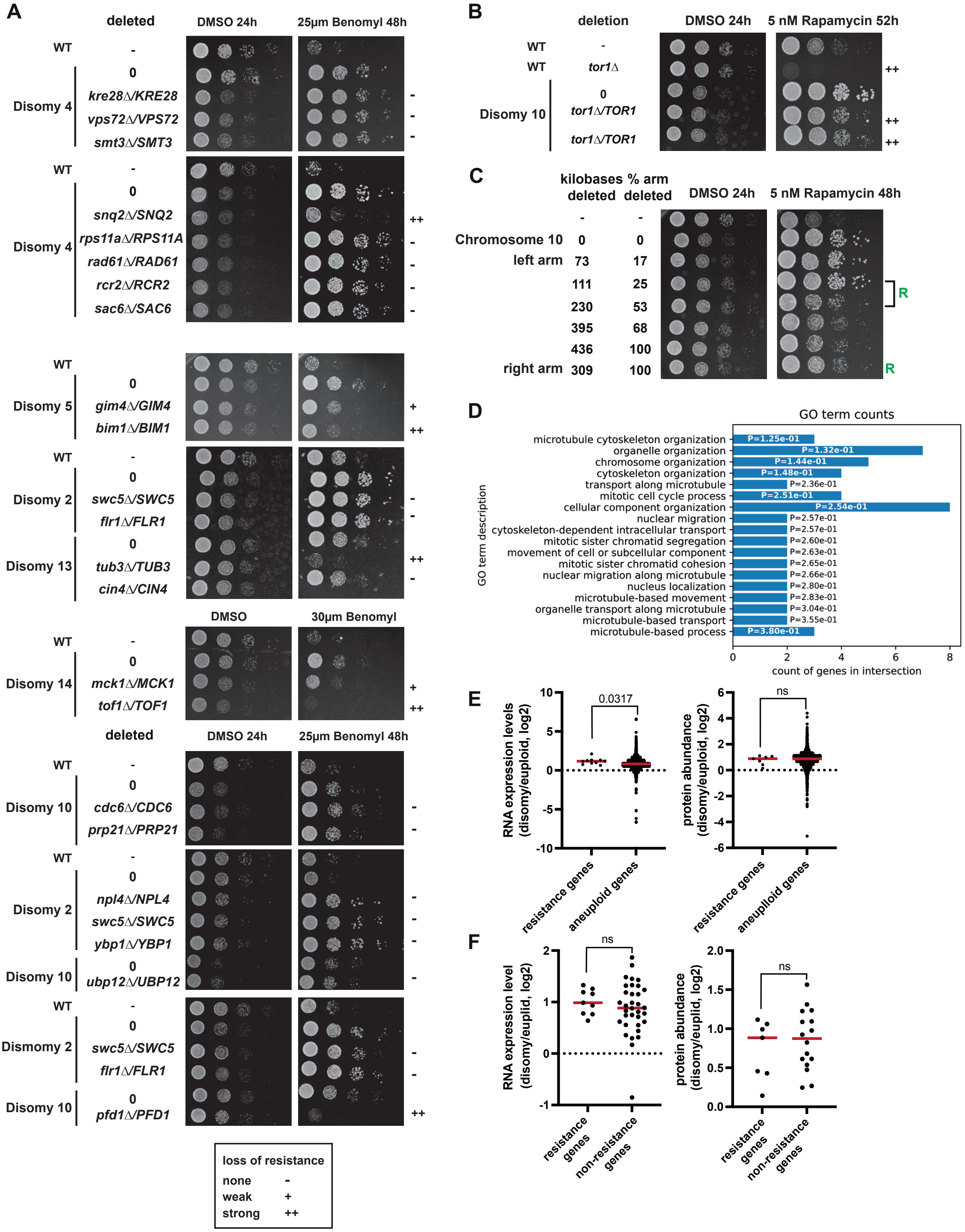

A
deleted
DMSO 24h
25μm Benomyl 48h
WT
Disomy 4
kre28Δ/KRE28
vps72Δ/VPS72
smt3Δ/SMT3
snq2Δ/SNQ2
rps11aΔ/RPS11A
rad61Δ/RAD61
rcr2Δ/RCR2
sac6Δ/SAC6
Disomy 5
gim4Δ/GIM4
bim1Δ/BIM1
Disomy 2
swc5Δ/SWC5
flr1Δ/FLR1
Disomy 13
tub3Δ/TUB3
cin4Δ/CIN4
DMSO
30μm Benomyl
Disomy 14
mck1Δ/MCK1
tof1Δ/TOF1
Disomy 10
cdc6Δ/CDC6
prp21Δ/PRP21
npl4Δ/NPL4
ybp1Δ/YBP1
ubp12Δ/UBP12
Dismomy 2
pfd1Δ/PFD1
loss of resistance
none -
weak +
strong ++
B
deletion
DMSO 24h
5 nM Rapamycin 52h
tor1Δ
tor1Δ/TOR1
C
kilobases deleted
% arm deleted
5 nM Rapamycin 48h
Chromosome 10
left arm
right arm
R
D
GO term counts
GO term description
microtubule cytoskeleton organization P=1.25e-01
organelle organization P=1.32e-01
chromosome organization P=1.44e-01
cytoskeleton organization P=1.48e-01
transport along microtubule P=2.36e-01
mitotic cell cycle process P=2.51e-01
cellular component organization P=2.54e-01
nuclear migration P=2.57e-01
cytoskeleton-dependent intracellular transport P=2.57e-01
mitotic sister chromatid segregation P=2.60e-01
movement of cell or subcellular component P=2.63e-01
mitotic sister chromatid cohesion P=2.65e-01
nuclear migration along microtubule P=2.66e-01
nucleus localization P=2.80e-01
microtubule-based movement P=2.83e-01
organelle transport along microtubule P=3.04e-01
microtubule-based transport P=3.55e-01
microtubule-based process P=3.80e-01
count of genes in intersection
E
RNA expression levels (disomy/euploid, log2)
0.0317
ns
protein abundance (disomy/euploid, log2)
resistance genes
aneuploid genes
F
RNA expression level (disomy/euplid, log2)
protein abundance (disomy/euplid, log2)
resistance genes
non-resistance genes

**Figure S7**

(A) 10-fold serial dilutions of candidate benomyl resistance genes. Phenotype classifications are annotated on the right and characterized as: none (-), weak (+), and strong (++). (B) 10-fold serial dilution of ∆*tor1* phenotypes on rapamycin. The phenotype classifications were annotated as for panel A. (C) 10-fold serial dilutions of partial chromosome arm deletions of chromosome 10. Control plates (YPAD + DMSO) were imaged after 24h, and treated plates (YPAD + 5nM rapamycin) after 48h. Regions associated with resistance are indicated with an (R) on the right side. (D) Gene ontology over-representation analysis for identified benomyl resistance driver genes. Absolute FDR adjusted p-values (mHG model and Benjamini and Hochberg correction) are shown. (E) Expression level increase on aneuploid chromosomes for benomyl resistance genes compared to all genes on the aneuploid chromosomes. Means are indicated by red lines, p-values are from Mann-Whitney U tests. (F) Expression levels of identified benomyl resistance genes compared to the expression levels of candidate genes that didn't contribute to benomyl resistance. Means are indicated by red lines, p-values are from Mann-Whitney U tests. Transcriptome data are from Torres et al. 2007 (chromosomes 2, 4, 5, 10, 13, 14). Proteome data are from Dephoure et al. 2014 (chromosomes 2, 5, 10, 13, 14).

#### Figure S8

| Chromosome | Region | Type of region | Gene |
|---|---|---|---|
| 2 | 0 - 237,423 | none | |
| Cen2 | - - - - - | - - - - - | - - - - - |
| | 238,979 - 341,042 | sensitivity | *FLR1* |
| | 341,042 - 436,902 | none | |
| | 436,902 - 537847 | none | |
| | 537,847 - 589,726 | resistance | *CDC28, SLI15, APD1, NPL4* |
| | 589,726 - 642,519 | none | |
| | 642,519 - 665,194 | resistance | *YBP1, AME1* |
| | 665,194 - 686,861 | none | *SWC5, DAD3* |
| | 686,861 - 713,057 | none | |
| | 713,057 - 737,873 | none | |
| 4 | 0 - 449,243 | sensitivity | |
| Cen4 | - - - - - | - - - - - | - - - - - |
| | 449-859 - 497,791 | resistance | ***SNQ2,*** *RCR2, RPS11A, RAD61* |
| | 497,791 - 598898 | none | |
| | 598,898 - 700,500 | resistance | *ARP10, PDS1, DPB4, APC4* |
| | 700,500 - 899,451 | none | *SAC6* |
| | 899,451 - 1,071,899 | none | *CHL4* |
| | 1,071,899 - 1,172,361 | resistance | ***SWR1*** |
| | 1,172,361 - 1,384,937 | none | |
| | 1,384,937 - 1,531,933 | sensitivity | *EFT2, SPC110, ERD1, SMT3, LCD1, VPS72, KRE28* |
| 5 | 0 - 76,019 | none | |
| | 76,019 - 133,182 | mild resistance | |
| | 133,182 - 151,182 | resistance | ***GIM4*** |
| Cen5 | - - - - - | - - - - - | - - - - - |
| | 152,907 - 182,637 | none | *PAC2* |
| | 182,637 - 242,199 | resistance | ***BIM1***, *GLN3* |
| | 242,199 - 301,923 | resistance | ***SAP1*** |
| | 301,923 - 364,508 | none | |
| | 364,508 - 415,848 | none | *MAM1* |
| | 415,848 - 576,874 | resistance | |
| 10 | 0 - 72,703 | mild resistance | *CDC6, PRP21, ECM25, UBP12* |
| | 72,703 - 110,725 | resistance | ***PFD1*** |
| | 110,725 - 220,012 | none | |
| | 220,012 - 328,956 | none | |
| | 328,956 - 435,981 | sensitivity | *MAD3, MAD2* |
| Cen10 | - - - - - | - - - - - | - - - - - |
| | 436,802 - 745,751 | none | |
| 12 | 0 - 150,711 | none | |
| Cen12 | - - - - - | - - - - - | - - - - - |
| | 151,725 - 489,957 | strong sensitvity | |
| | 489,957 - 1,078,177 | sensitivity | *TUB4* |
| 13 | 0 - 268,031 | resistance | ***TUB3***, *TUB1* |
| Cen13 | - - - - - | - - - - - | - - - - - |
| | 268,149 - 924,431 | none | *CIN4* |
| 14 | 0 - 120,998 | mild resistance | ***MCK1***, *CLA4* |
| | 120,998 - 220,164 | resistance | ***TOF1*** |
| | 220,164 - 320,127 | none | |
| | 320,127 - 627,772 | mild sensitivity | *ALF1* |
| Cen14 | - - - - - | - - - - - | - - - - - |
| | 629,032 - 784,333 | none | |

**Figure S8**

Phenotypes for regions between all full and partial chromosome arm deletions engineered in this study. Centromeres are indicated by a dashed line. Phenotypes that were assigned to a region are annotated in 'Type of region'. Three types of phenotypes were assigned: none – no change in phenotype upon deletion, sensitivity – increased benomyl resistance upon deletion, mild sensitivity – mild decrease in benomyl resistance upon deletion, resistance – substantial decrease in in benomyl resistance upon deletion. All genes tested are listed; genes that decrease resistance when heterozygously deleted in the aneuploid strain are in bold.

#### Figure S9

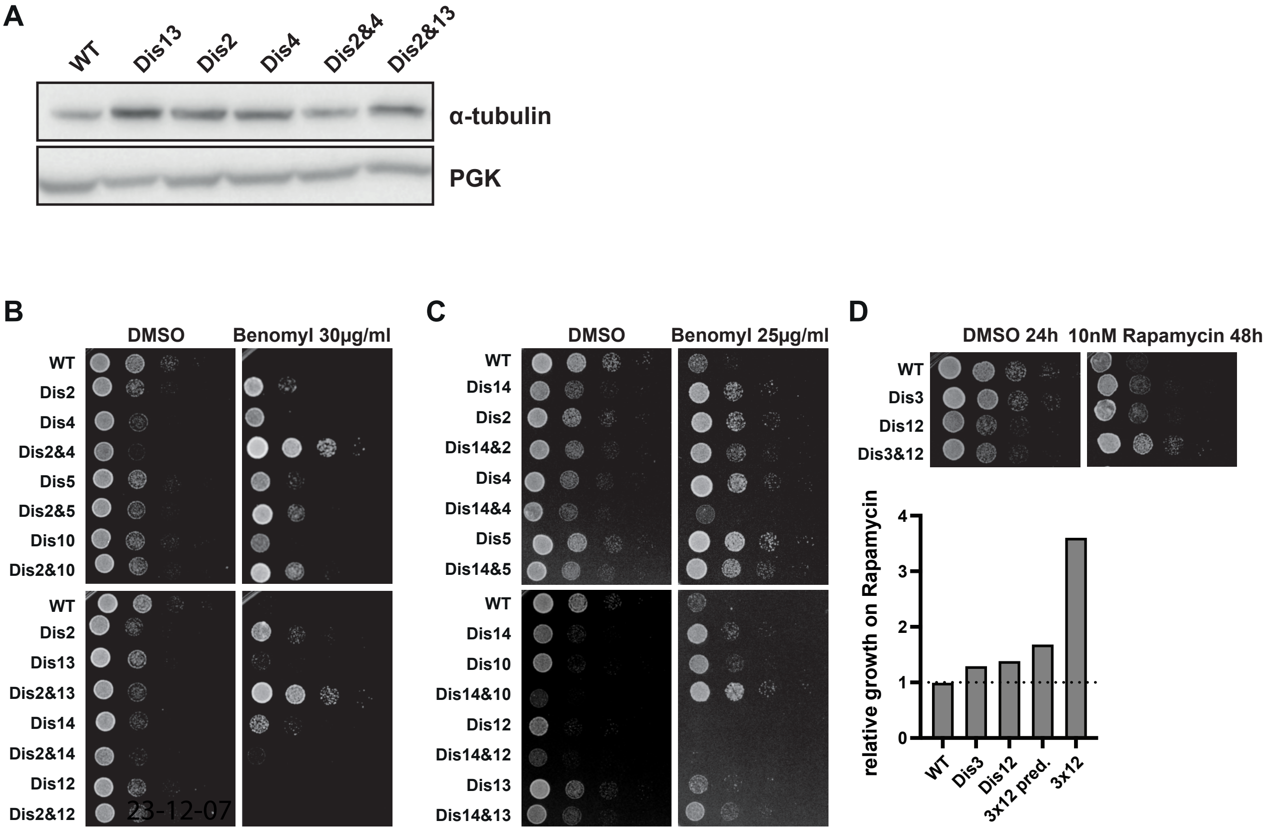


**Figure S9**
(A) Western blot analysis of α-tubulin expression levels for the indicated single and double disomies. (B) 10-fold serial dilutions of disomy combinations with chromosome 2. Control plates (YPAD + DMSO) were imaged after 24h, and treated plates (YPAD + 30 µg/ml benomyl) after 48h. (C) 10-fold serial dilutions of disomy combinations with chromosome 14. Control plates (YPAD + DMSO) were imaged after 24h, and treated plates (YPAD + 25 µg/ml benomyl) after 48h. (D) 10-fold serial dilutions and quantifications of disomy 3 and 12 on control (YPAD + DMSO) and treated (10 nM rapamycin) plates. All quantifications were normalized to a haploid WT. Combinatorial growth of chromosome 3 and 12 double disomy was predicted by adding the phenotypes of the single disomies.
